# Manganese scavenging is required for virulence of the cytosolic pathogen *Listeria monocytogenes*

**DOI:** 10.64898/2026.10.06.757198

**Authors:** Carly A. Mascari, Christal Carerros, Jessica L. Kelliher

## Abstract

Metals are essential for all forms of life and are therefore the subject of tight competition at the host-pathogen interface. Manganese (Mn) is an important cofactor for enzymes involved in various processes in bacteria, including the response to oxidative stress, and the ability to scavenge Mn through the expression of high-affinity Mn transporters has been associated with virulence of many extracellular and vacuolar pathogens. While the cytosol has been established to be restrictive for growth of non-adapted pathogens, the extent to which metal limitation contributes to this is not well understood. Here, we investigate whether high-affinity Mn acquisition is important for intracellular adaptation and virulence of the cytosolic pathogen *Listeria monocytogenes* (*Lmo*). To do so, we generated a strain of *Lmo* in which both of its predicted high-affinity Mn transporters, MntH and MntABC, are disrupted. We found that this strain was severely defective in growth *in vitro* under Mn-limited conditions and had lower intracellular Mn content. Additionally, we observed Mn-dependent transcriptional regulation of the *mntH* and *mntA* genes. Together, these data strongly support that MntH and MntABC are high-affinity Mn transporters and are likely to be the only such dedicated Mn transporters expressed in *Lmo.* Using the Δ*mntH* Δ*mntA* strain, we provide evidence that Mn limitation sensitizes *Lmo* to oxidative stress *in vitro*, consistent with the known role of Mn in mitigating reactive oxygen species. Ultimately, we demonstrate that expression of high-affinity Mn transporters is essential for intracellular replication of *Lmo* in macrophages, plaquing on fibroblast monolayers, and for virulence in an animal model of systemic infection. Our work, therefore, suggests that *Lmo* is challenged by Mn limitation in the cytosol and requires Mn scavenging to adapt to its intracellular niche.

**Author summary:** Both bacterial pathogens and their hosts require metals, such as manganese (Mn), to survive, and the mammalian host has evolved to exploit this necessity by restricting metal availability at the host-pathogen interface. To overcome this, pathogens typically express high-affinity metal transport systems to allow them to compete for metals. Mn-scavenging transporters are essential for virulence of a number of pathogens that replicate in the extracellular space and intracellularly within vacuoles. However, for pathogens such as *Listeria monocytogenes* (*Lmo*) that can replicate within the cytosol of host cells, whether Mn is limited within the cytosol and whether these pathogens require Mn scavenging to survive in this niche has been less thoroughly investigated. Here, we establish the transporters used by *Lmo* for Mn scavenging and demonstrate that without these systems, this pathogen is defective in intracellular replication in cells *ex vivo* and virulence in an animal model. Thus, our results indicate that *Lmo* is challenged by Mn limitation in the cytosol and requires Mn scavenging systems to adapt to this environment.

## Introduction

All organisms must obtain inorganic nutrients from their environment. This includes transition metals, such as iron (Fe), zinc (Zn), and manganese (Mn), which act as cofactors for the structure or function of an estimated one-third to one-half of all proteins (1, 2). At the host-pathogen interface, vertebrates obtain these essential nutrients from dietary sources, while pathogenic microorganisms acquire metals directly from the host. This creates a tug-of-war dynamic wherein host cells and pathogens are in direct competition for these often-limiting nutrients. This competition has led to the evolution of host defense strategies, collectively referred to as nutritional immunity, in which the host actively restricts metal availability in response to infection to starve invading pathogens (3). In turn, pathogenic microorganisms have evolved mechanisms to respond to nutrient limitation, such as scavenging metals via high-affinity metal acquisition systems (3, 4).

Mn is an important nutrient with diverse functions in bacteria, including in glycolysis, signal transduction, and nucleotide metabolism (5). Mn also plays an important role in oxidative stress resistance by acting as a cofactor for a subset of superoxide dismutase (SOD) enzymes (6) and by replacing Fe in mononuclear enzymes to reduce the cellular reliance on Fe, which, unlike Mn, participates in Fenton chemistry (7, 8). In accordance with the importance of Mn for bacteria, the sequestration or mobilization of Mn has been shown to be a powerful antimicrobial defense strategy employed by the mammalian host. For example, secretion of the metal-binding S100 family protein calprotectin (CP) by neutrophils at sites of *Staphylococcus aureus* abscesses in the liver results in localized depletion of Mn and limits bacterial burdens during infection (9, 10). While *S. aureus* lacking the ability to scavenge Mn is less virulent during systemic infection, this phenotype is reversed in the Mn-replete liver of CP^−/−^ mice (11), highlighting the tight competition for Mn at this infectious interface. Another notable example is the mobilization of Mn and other divalent metals out of the phagolysosome by the host-encoded transporter Natural resistance-associated macrophage protein 1 (Nramp1) (3). Inbred mouse lines with loss-of-function mutations in the gene encoding Nramp1 (including the commonly used C57BL/6 and BALB/c lines) are more susceptible to various vacuolar pathogens (12–14) due to an inability to deplete the vacuole of divalent metals. Notably, while these examples highlight the dynamic nature of Mn at the host-pathogen interface in the extracellular space and within vacuoles, considerably less is known about Mn dynamics within the mammalian cytosol.

A conserved mechanism for dealing with metal limitation across bacteria is the expression of high-affinity metal acquisition systems. While non-cognate transporters can be permissive for uptake in metal-replete environments, dedicated transporters become necessary for high-affinity scavenging when the substrate is in low abundance. For Mn, the two most common transporter types in bacteria are the Nramp1 family Mn^2+^/H^+^ symporters, MntH, and the ATP-binding cassette Mn^2+^ transporters, MntABC (5, 15). MntABC homologs consist of a substrate-binding lipoprotein, a transmembrane permease protein, and a cytoplasmic ATP-binding protein. Many bacterial species encode at least one copy of both transporter types, though the distinct roles of the two transporters are not yet well understood. Differences in affinity for Mn or activity under certain growth conditions have been reported in some organisms (5). Notably, MntH homologs have been proposed to be more active at acidic pH, while MntABC homologs may function optimally at neutral or slightly alkaline pH (16–19). In any case, expression of one, or either, type of high-affinity Mn importer is required for virulence of a number of pathogens, including *Staphylococcus aureus* (11, 20), *Streptococcus pneumoniae* (21, 22)*, Salmonella* Typhimurium (23)*, Enterococcus faecalis* (24, 25)*, Bacillus anthracis* (26) and *Yersinia pestis* (27), highlighting the importance of Mn scavenging in the host.

*Listeria monocytogenes* (*Lmo*) is a clinically significant, cytosolic pathogen that comprises 18% of the total economic burden from foodborne pathogens in the United States (28). In immunocompromised and pregnant individuals, infection with *Lmo* (called listeriosis), can result in septicemia, central nervous system infection, and fetal demise. Upon ingestion by the host via contaminated food products, *Lmo* promotes its own internalization into intestinal epithelial cells. Once internalized, *Lmo* uses the virulence factor listeriolysin O to perforate the vacuole and escape into its replicative niche, the cytosol. In the cytosol, *Lmo* hijacks host actin, allowing it to project through the cell and ultimately spread directly cell-to-cell where this cycle repeats. *Lmo* is also capable of escaping phagocytic vacuoles and replicating rapidly within the cytosol of professional phagocytes like macrophages (29). Ultimately, the ability of *Lmo* to thrive within the cytosol of both phagocytic and non-phagocytic cells is essential for its pathogenesis. The cytosol is restrictive to non-adapted microbes through sparsely understood cell-intrinsic defenses and nutritional constraints (30–32), and we and others have identified specific adaptations required by *Lmo* to survive and replicate in this niche (33–41). These include the ability scavenge essential nutrients like lipoic acid and riboflavin-derived cofactors, for which *Lmo* is auxotrophic (40, 41). Expression of dedicated Fe and Zn transporters has also been found to be critical for *Lmo* to replicate intracellularly (38) and cause systemic infection (38, 39), together suggesting that nutrient limitation is a significant challenge faced by *Lmo* in the cytosol.

Previous studies have established that Mn is an essential nutrient for *Lmo in vitro* (42, 43) and have identified putative MntH and MntABC homologs in the listerial genome (18, 44, 45). A mutant expressing a truncated version of *mntH* has a severe growth defect in acidic but not neutral pH conditions, a defect that could be rescued by addition of excess Mn, suggesting that MntH is a *bona fide* Mn importer (18). The role of MntH in supporting the intracellular lifestyle and virulence of *Lmo* has not been directly addressed. The *mntABC* locus was first studied in *Lmo* due to homology of the *mntA* (metal-binding lipoprotein) gene product to the streptococcal protein PsaA (44). In addition to its role in Mn import, PsaA facilitates adhesion of *Streptococcus* species to host cells (46). While no differences in adhesion were observed for an *Lmo mntA* deletion mutant, loss of MntA was found to be associated with reduced invasion in epithelial and hepatocyte cell lines (44, 47) and reduced plaque number in epithelial cells (47). However, loss of *mntA* did not impair bacterial replication within macrophages or significantly reduce virulence in either an intravenous mouse (44) or a *Galleria mellonella* infection model (47). Given that these earlier studies focused on the possible adhesin role of MntA, the role of MntABC in Mn transport in *Lmo* and the contribution of Mn acquisition to the observed phenotypes of the *mntA* deletion strain has not yet been addressed. Interestingly, both *mntH* and *mntABC* transcripts were found to be upregulated in *Lmo* isolated from the spleens of infected mice (48), which may be consistent with *Lmo* experiencing Mn limitation *in vivo*. However, the extent to which *Lmo* is challenged by Mn starvation in the host, including in its cytosolic niche, is not clear.

In order to address whether Mn acquisition is essential for intracellular adaptation and virulence, we sought to disrupt the high-affinity Mn importers in *Lmo* to diminish Mn scavenging. After generating a Δ*mntH* Δ*mntA* double mutant strain, we validated that this mutant was impaired in Mn acquisition and growth under Mn-limiting conditions. We then utilized this strain to evaluate the importance of Mn scavenging for reactive oxygen species stress response, cytosolic adaptation, and virulence of *Lmo*. Ultimately, we found that the ability of this cytosolic pathogen to acquire Mn is critical for its survival in the cytosol and capacity to cause invasive infection.

## Results

### An *Lmo* strain lacking both *mntH* and *mntA* is severely diminished in Mn scavenging

To determine the extent to which Mn scavenging contributes to the pathogenesis of *Lmo*, we first set out to identify its dedicated high-affinity Mn acquisition systems. Previous studies had identified a single copy each of *mntH* and *mntABC* in the *Lmo* genome (18, 44, 45). To rigorously evaluate the role of these loci in Mn scavenging *in vitro*, we generated in-frame deletion strains lacking either *mntH* or *mntA* (the substrate-binding component of the MntABC transporter), or both, in the 10403S background. While Δ*mntH* and Δ*mntA* single mutants were readily recovered on Brain Heart Infusion (BHI) agar following allelic exchange, we were unable to generate the Δ*mntH* Δ*mntA* strain. However, supplementation of BHI agar with 50 µM Mn (MnSO_4_) allowed us to recover the double mutant strain at the expected ∼50% frequency. To validate this observation, we subsequently grew the Δ*mntH* Δ*mntA* strain in parallel with both single mutant strains and wild type in Mn-replete medium overnight, then replica spotted on BHI with or without Mn supplementation. The Δ*mntH* Δ*mntA* strain was effectively unable to grow on standard BHI agar, but growth was completely rescued in the presence of 50 µM supplemental Mn (**Fig 1A**). Neither single mutant strain was growth-impaired on BHI agar in this assay, consistent with prior studies showing that disruption of either the *mntH* or *mntABC* locus individually is not sufficient to impact growth of *Lmo* in rich medium (18, 44).

**Fig 1.**
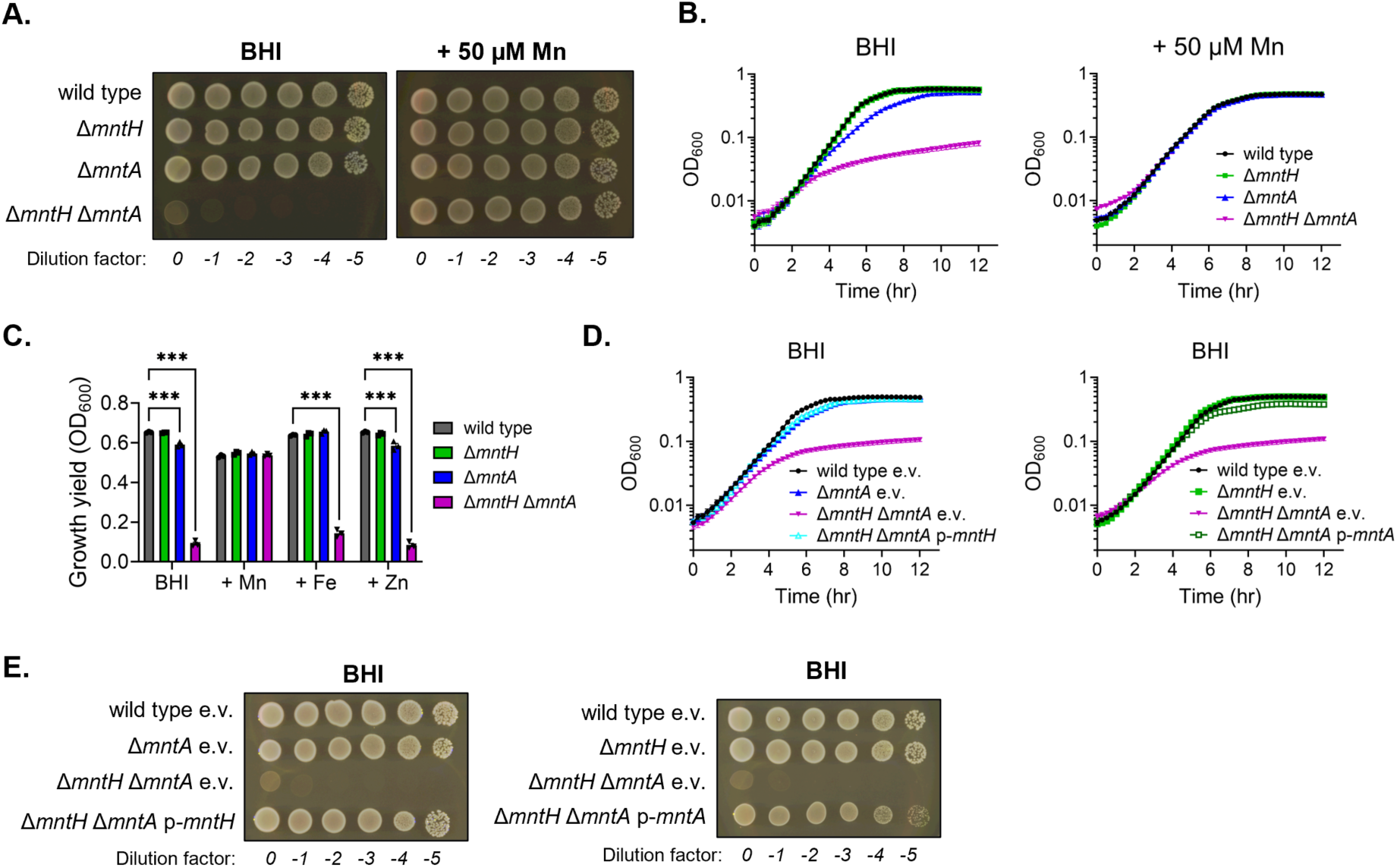
Simultaneous disruption of MntH and MntABC impairs growth of *Lmo* in rich medium in a Mn-dependent manner. (A) and (E) Spot dilution assay for the indicated strains of *Lmo* grown overnight at 37°C on standard BHI agar or BHI agar supplemented with 50 µM Mn as indicated. Images are representative of three independent experiments. (B) and (D) Growth curves for the indicated strains in BHI liquid culture ±50 µM Mn as indicated. OD_600_ measurements were recorded every 15 min. (C) Growth yields (OD_600_) at 12 hr for the indicated strains grown in BHI supplemented, or not, with 50 µM Mn, Fe or Zn. Also see **S1B Fig** for full growth curves. (B-D) Error bars represent SEM; n = 3 independent experiments each with technical triplicate measurements. (C) Statistical differences relative to wild type are shown and were determined by two-way ANOVA with Tukey’s multiple comparisons. *** = p ≤ 0.001. e.v. indicates empty vector.

We observed a similar, severe growth defect for the Δ*mntH* Δ*mntA* strain in BHI broth that, as with solid agar, could be rescued with Mn supplementation in a dose-dependent manner (**Fig 1B and S1A Fig**). Importantly, only Mn and not Fe or Zn supplementation resulted in wild type-like growth in the Δ*mntH* Δ*mntA* strain, though Fe supplementation did modestly improve growth relative to standard BHI (**Fig 1C** and **S1B Fig**). Supplementation of BHI with MnCl_2_ instead of MnSO_4_ also fully rescued growth of the Δ*mntH* Δ*mntA* strain, suggesting that Mn itself, and not the counterion, is driving this phenotype (**S1C Fig**). Additionally, the growth defect of the Δ*mntH* Δ*mntA* strain in both BHI liquid culture and on BHI agar could be genetically complemented by expression *in trans* of either *mntH* or *mntA* from integrative vectors (**Fig 1D-E**).

Interestingly, the *mntA* deletion strain displayed a minor growth defect in liquid culture in BHI (**Fig 1B**) that could be genetically complemented *in trans* (**S1D Fig**), but there was no growth impairment for the Δ*mntH* strain. Supplementation of BHI with either Mn or Fe (but not Zn) resulted in wild type-like growth for the Δ*mntA* strain (**Fig 1C and S1B Fig**). We confirmed that strains in which the entire *mntABC* operon was deleted rather than only *mntA*, *i.e.*, Δ*mntABC* and Δ*mntH* Δ*mntABC* mutants, grew similarly to the Δ*mntA* and Δ*mntH* Δ*mntA* mutants, respectively, on BHI agar and in broth (**S1E**-**F Fig**). Taken together, this consistent with MntA functioning as part of an MntABC complex which may have some promiscuity for Fe import.

Given that BHI is a complex medium in which metal levels cannot be easily manipulated, and that growth yields were slightly reduced in BHI supplemented with Mn relative to standard BHI for wild type and the single mutant strains (**Fig 1C**), we employed the defined *Listeria* Synthetic Medium (LSM) (43) to enable control over Mn concentration. The standard LSM recipe contains 50 µM MnSO_4_ (heretofore referred to as LSM-Mn50). In LSM-Mn50, all strains grew comparably in broth culture (**Fig 2A**); thus, we subsequently used LSM-Mn50 as our standard Mn-replete growth condition. Upon decreasing the Mn concentration in LSM, we observed a dose-dependent attenuation of growth for all strains (**S2A**-**B Fig**). However, this effect was greatly exacerbated in the Δ*mntH* Δ*mntA* strain. At 0.1 µM Mn, growth rate of the Δ*mntH* Δ*mntA* mutant began to slow relative to wild type during mid-exponential phase, suggesting that Mn was becoming limiting to this strain (**S2A Fig**). Thus, LSM containing 0.1 µM Mn (LSM-Mn0.1) was selected as our Mn-limiting condition for experiments going forward (**Fig 2A**).

**Fig 2.**
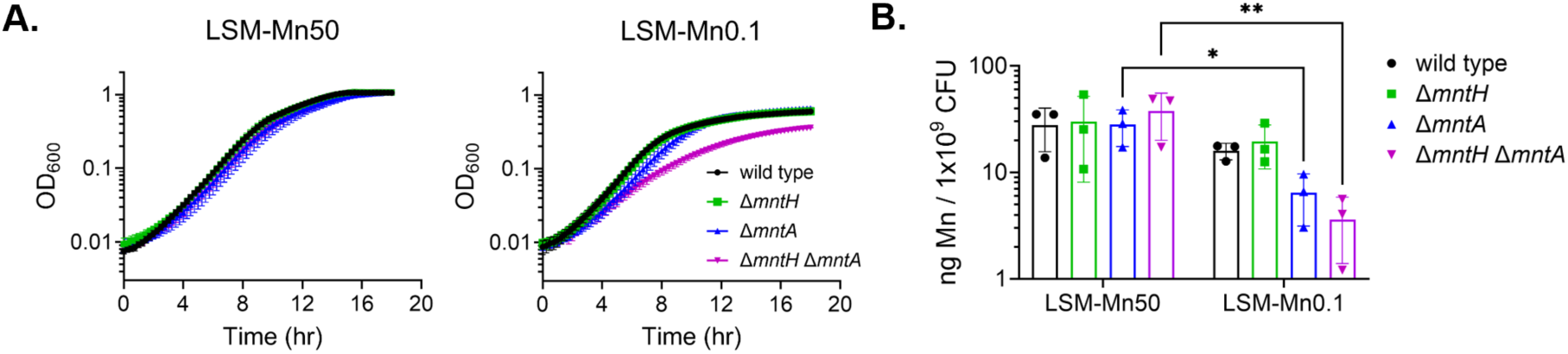
A Δ*mntH* Δ*mntA* strain is growth-impaired in Mn-limited minimal medium and exhibits reduced intracellular Mn. (A) Growth curves for the indicated strains in liquid culture in *Listeria* Synthetic Medium (LSM) containing either 50 µM (left) or 0.1 µM (right) Mn. OD_600_ measurements were recorded every 15 min. Error bars represent SEM; n = 3 independent experiments each with technical triplicate measurements. (B) Intracellular Mn levels in the indicated strains grown to mid-exponential phase in LSM-Mn50 or LSM-Mn0.1 as determined by inductively coupled plasma mass spectrometry (ICP-MS) and normalized to colony-forming units (CFU). Error bars represent SD; n = 3 independent experiments. Statistical differences were determined by two-way ANOVA with Tukey’s multiple comparisons. * = p < 0.05; ** = p < 0.01.

To further validate the role of MntH and MntABC in Mn scavenging, we utilized inductively coupled plasma mass spectrometry (ICP-MS) to measure intracellular metal levels during growth in Mn-replete and Mn-limited conditions. Wild type and all three mutant strains were grown to mid-exponential phase in LSM-Mn50 and LSM-Mn0.1. Under these growth conditions, both wild type and the Δ*mntH* Δ*mntA* strain grew similarly, with only minor growth impairment observed for the Δ*mntH* Δ*mntA* in LSM-Mn0.1 relative to LSM-Mn50 as mentioned above (**S2C Fig**). We observed a decrease in intracellular Mn for all strains when grown in LSM-Mn0.1 compared to LSM-Mn50; however, this difference was statistically significant for the Δ*mntA* single mutant and most pronounced for Δ*mntH* Δ*mntA* double mutant strain (**Fig 2B**). This is consistent with a model in which MntABC plays a more critical role than MntH in Mn acquisition under these growth conditions and indicates that the Δ*mntH* Δ*mntA* strain is highly impaired in Mn scavenging.

Mn transporters have been found to have some promiscuity for import of other divalent metals, including Zn and Fe (15). We did not observe any differences in intracellular Zn content between strain when grown in standard LSM-Mn50 and only observed a slight increase in intracellular Zn in the *mntH* deletion strain when grown in LSM-Mn0.1 relative to LSM-Mn50 (**S2D Fig**). We did, however, observe a significant increase in intracellular Fe for both wild type and the *mntH* deletion strain, but not Δ*mntA* or Δ*mntH* Δ*mntA*, under Mn-limited conditions (LSM-Mn0.1, **S2E Fig**). This observation points to an MntABC-dependent increase in Fe import when Mn is limited, though it is not clear if this is occurring directly via the MntABC transporter itself, or as a regulatory response to loss of Mn-acquisition via MntABC. That said, we found that both wild-type and Mn transporter mutant strains of *Lmo* were largely insensitive to the Fe content of LSM both under Mn-replete and Mn-limited conditions (**S2F**-**G Fig**). In other words, the growth defect of the double transporter mutant in BHI and LSM-Mn0.1 is not due to a reduced ability to acquire Fe. Thus, while it remains possible that MntABC is able to import Fe in addition to Mn, this does not appear to be its primary physiological role at least under the conditions tested. Instead, our findings indicate that a *mntH mntA* double mutant strain is significantly impaired in Mn scavenging such that growth of this strain is impaired when Mn availability is limited.

### *mntH* and *mntABC* are regulated in a Mn-dependent manner

MntH and MntABC homologs in other species are regulated in a Mn-dependent manner wherein transporter expression is repressed when Mn is abundant to prevent uncontrolled import and subsequent Mn toxicity (20, 49–53). When Mn levels become limiting, expression is de-repressed to permit Mn scavenging. To evaluate Mn transporter regulation in *Lmo,* we performed RT-qPCR on lysates from wild-type *Lmo* grown at high (50 µM), intermediate (1 µM), or low (0.1 µM) Mn concentration in LSM using primers specific to either *mntH* or *mntA.* While we observed little to no induction of either transcript at 1 µM relative to 50 µM Mn, growth at 0.1 µM Mn resulted in ∼12- and ∼60-fold induction of *mntH* and *mntA*, respectively, relative to Mn-replete conditions (**Fig 3A**-**B**). The observation that expression of both transporters is increased in Mn-limiting growth conditions is consistent with the idea that they have a role in Mn scavenging.

**Fig 3.**
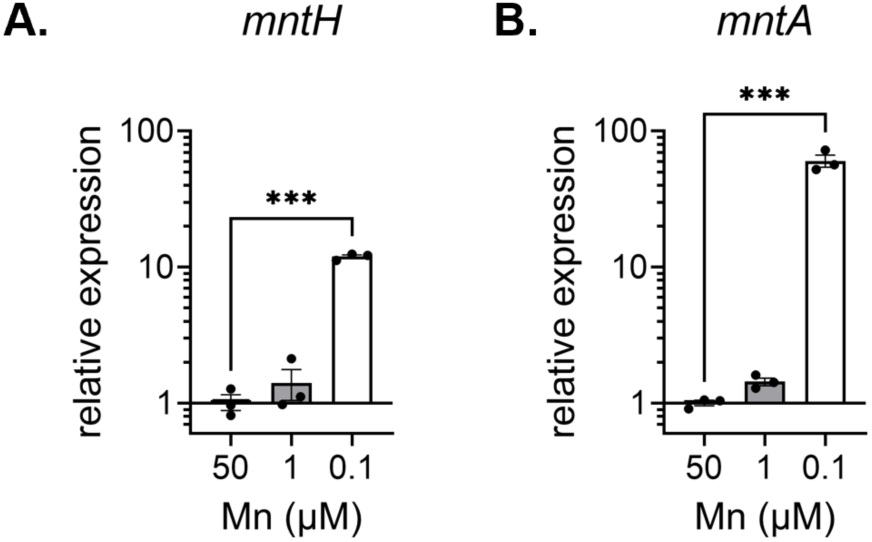
Expression of *mntH* and *mntA* is increased during Mn limitation. (A) *mntH* and (B) *mntA* transcripts from cells grown to mid-exponential phase in LSM containing either 50, 1, or 0.1 µM Mn were quantified using RT-qPCR and analyzed by the Pfaffl method. *16s rRNA* was used as the reference gene and expression was reported relative to growth at 50 µM Mn. Error bars represent SEM; n = 3 independent replicates measured in technical triplicate. Statistically significant differences relative to 50 µM Mn are shown and were determined by one-way ANOVA with Tukey’s multiple comparisons. *** = p ≤ 0.001.

### Disruption of Mn scavenging impairs ROS mitigation *in vitro*

*Lmo* encodes one constitutively expressed, Mn-dependent SOD (54–56). This, taken together with the role of Mn in Fe replacement (7, 8), led us to hypothesize that the growth defect of the Δ*mntH* Δ*mntA* strain under Mn-limited conditions may be in part due to diminished oxidative stress resistance. We reasoned if this is true, growth of the Δ*mntH* Δ*mntA* strain may be partially rescued in Mn-limiting conditions by being grown anaerobically. After growing aerobically in LSM-Mn50 overnight, the single and double transporter mutant strains were replica spotted on BHI agar with and without supplemental Mn, then incubated either exposed to air or anaerobically. While we observed less growth overall for all strains under anaerobic conditions, growth of the double mutant strain was more similar to wild type under anaerobic relative to aerobic conditions (**Fig 4A**). The fact that growth of the Δ*mntH* Δ*mntA* strain was not fully rescued by removal of oxygen suggests that while endogenous ROS stress is a contributor to the growth defect in this strain, it is, unsurprisingly, not the only driving factor.

**Fig 4.**
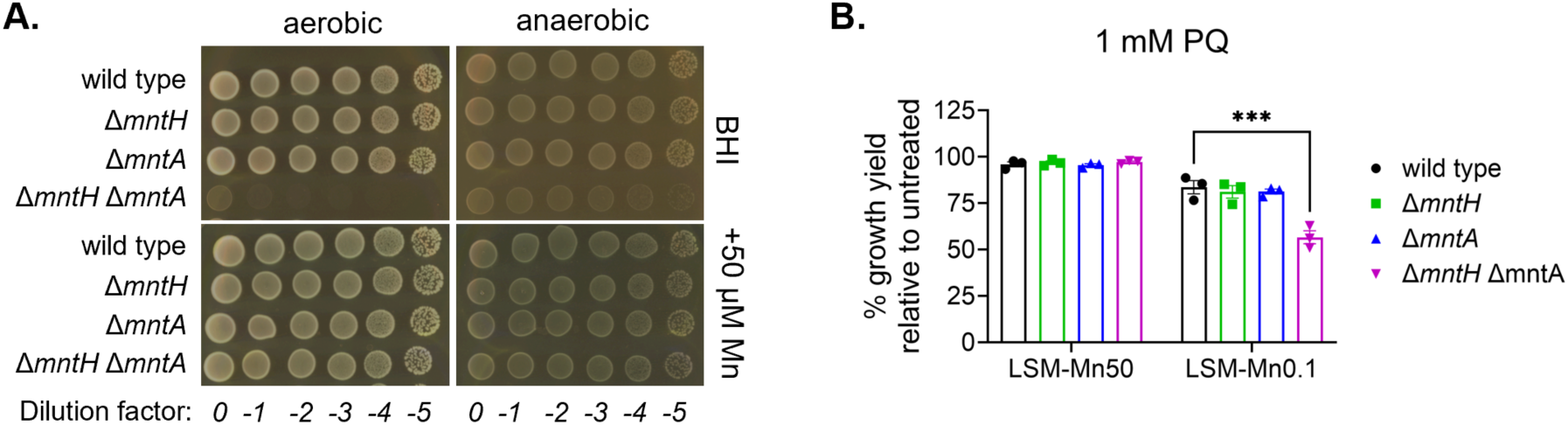
The Δ*mntH* Δ*mntA* strain is sensitized to aerobic growth and superoxide stress. (A) Spot-dilution assay for the indicated strains on BHI agar (top) or BHI agar supplemented with 50 µM Mn (bottom) grown overnight either exposed to air (left) or in an anaerobic pouch (right). Images are representative of two independent experiments. (B) *Lmo* strains were grown in the presence or absence of 1 mM paraquat (PQ) in LSM-Mn50 and LSM-Mn0.1. Final growth yield in the presence of PQ at 18 hr, as determined by OD_600_, is reported for each strain as a percentage of growth yield in the absence of PQ. Error bars represent SEM; n = 3 independent experiments. Statistical differences compared to wild type within each treatment are shown and were determined by two-way ANOVA with Tukey’s multiple comparisons. *** = p ≤ 0.001.

To further evaluate the extent to which Mn scavenging is critical for ROS mitigation, we tested sensitivity of the Δ*mntH* Δ*mntA* and single mutant strains to paraquat (PQ), a quaternary ammonium compound that generates intracellular superoxide radicals (57). Treatment with 1 mM PQ largely did not impact final growth yield in wild type and the single and double transporter mutant strains when grown in LSM-Mn50. At low Mn concentration (LSM-Mn0.1), PQ treatment reduced the final growth yield ∼20% for the wild-type and single mutant strains; however, the growth impairment of the double mutant strain was significantly increased compared to wild type, at ∼45% reduction relative to untreated (**Fig 4B, S3A Fig**), consistent with increased sensitivity of this strain to superoxide radicals. To determine whether the increased susceptibility of the Δ*mntH* Δ*mntA* mutant to PQ stems from diminished metalation of its Mn-dependent SOD, we evaluated SOD activity in lysates of cells grown in LSM-Mn50 and LSM-Mn0.1 supplemented, or not, with 1 mM PQ to induce superoxide stress. Cells were harvested at mid-exponential phase (**S2C Fig**), normalized for total protein content, and SOD activity was determined. While we did detect trending decreases in SOD activity in the Δ*mntH* Δ*mntA* strain in LSM-Mn0.1 relative to wild type both in the presence and absence of PQ, and increased SOD activity with PQ treatment for both strains, all differences observed were modest (**S3B Fig**). This result could suggest that under the growth conditions used here *Lmo* may not yet be Mn-limited enough to result in differences in SOD metalation and therefore activity.

Alternatively, given that Mn can mitigate oxidative stress through non-SOD mechanisms such as replacing Fe in mononuclear enzymes (7, 8), lack of such activities could account for the increased susceptibility to PQ of the Δ*mntH* Δ*mntA* strain. Regardless, these data cumulatively suggest that Mn acquisition by MntH and MntABC plays a role in resisting oxidative stress *in vitro* in *Lmo*.

### Mn scavenging is required for intracellular replication of *Lmo*

During infection, *Lmo* replicates within the host cell cytosol after escaping from either the endocytic or phagocytic vacuole. Adaptation to this hostile niche is essential for virulence of *Lmo* (35). To assess the importance of Mn scavenging for the intracellular lifecycle of *Lmo*, we first evaluated replication of wild type and our single and double Mn transporter mutant strains within bone marrow-derived macrophages (BMDMs) from C57BL/6 mice. Neither of the single mutant strains were impaired in intracellular replication, suggesting either transporter is sufficient to facilitate Mn acquisition intracellularly. In contrast, growth of the double transporter mutant was significantly diminished, reaching a lower burden than wild type at 5 hr post-infection, then ceasing replication thereafter (**Fig 5A**). This observed growth defect of the double mutant strain in macrophages could be genetically complemented by expression of either *mntH* or *mntA in trans* (**Fig 5B**-**C**). We could also significantly improve intracellular growth of the Δ*mntH* Δ*mntA* double mutant by supplementation of the culture medium with 50 µM Mn, which did not affect growth kinetics of wild-type *Lmo* (**Fig 5D**). Together, these data suggest that either MntH or MntABC is required for intramacrophage replication of *Lmo* due to their role in Mn scavenging.

**Fig 5.**
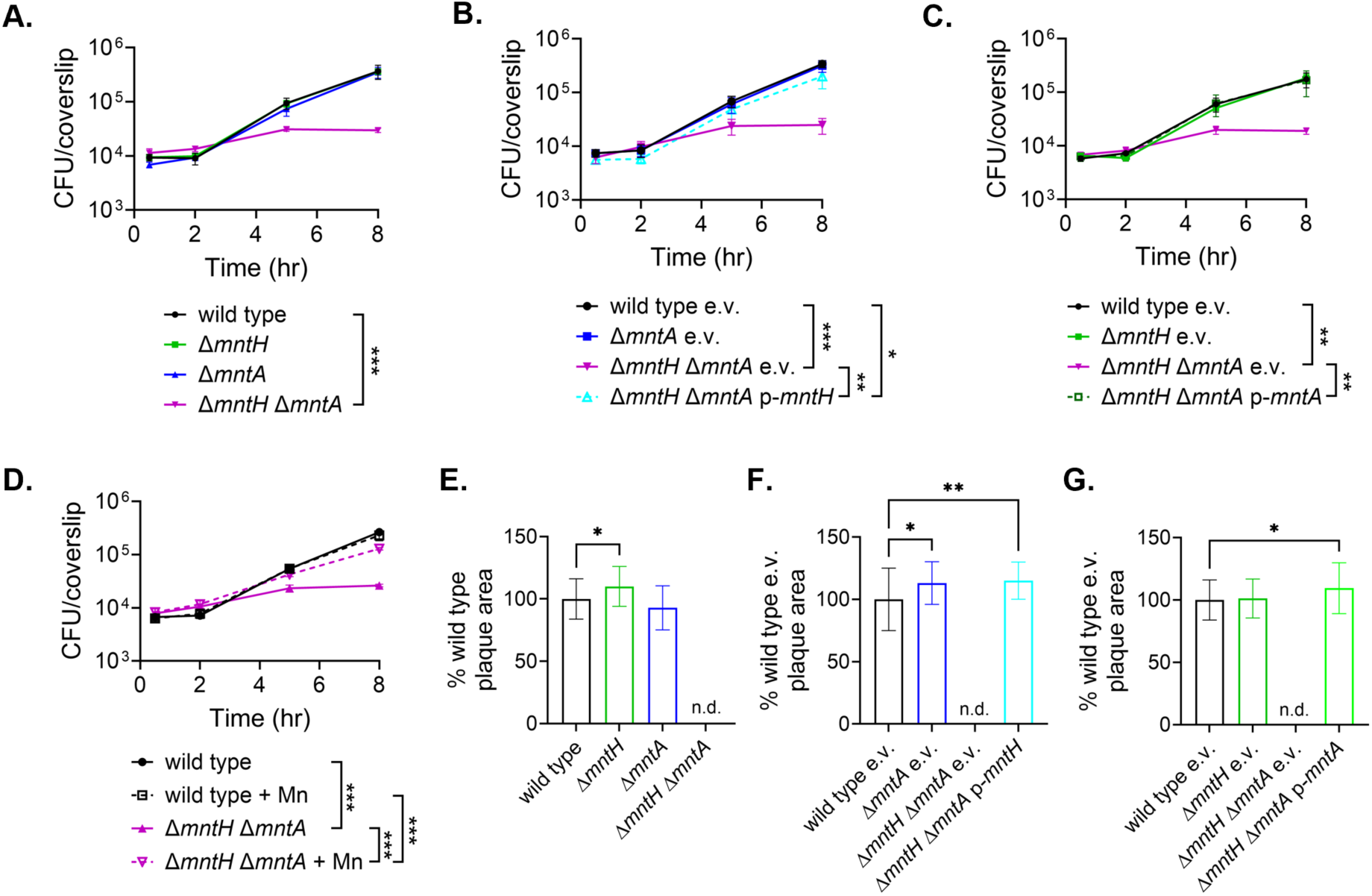
*Lmo* requires Mn scavenging for intracellular replication and virulence *ex vivo*. (A-D) Intracellular growth curves of the indicated strains within BMDMs. BMDMs were seeded on glass coverslips and CFUs were enumerated and averaged from three replicate coverslips at each timepoint. (D) 50 µM Mn was added, or not, to the cell culture medium at the time of BMDM seeding and maintained throughout the course of the experiment for the indicated strains. Error bars represent SEM; n = 3 independent experiments with technical triplicate measurements. Statistically significant differences were determined by two-way ANOVA with Tukey’s multiple comparisons. Asterisks indicate relevant differences at 8 hpi. (E-G) Plaque formation in immortalized murine fibroblasts (L2 cells). Plaque area was measured at 72 hpi across three independent experiments and is reported as a percentage of the average wild-type plaque area. Error bars represent mean ±SD. Statistical differences were determined by one-way ANOVA with Tukey’s multiple comparisons. Asterisks indicate differences relative to wild type. * = p < 0.05; ** = p < 0.01, *** = p ≤ 0.001. n.d. (not determined) indicates that no plaques were observed and e.v. indicates empty vector.

Given that we observed that *Lmo* deficient in Mn scavenging is sensitive to ROS *in vitro* (**Fig 4**) and that ROS production is a major host defense mechanism, we tested the hypothesis that the intracellular growth defect of the Δ*mntH* Δ*mntA* strain is due to increased susceptibility to host-derived ROS. We attempted to pharmacologically mitigate ROS during infection in BMDMs by treating cells at the time of infection with either the general ROS scavenger N-acetylcysteine (NAC) or the mitochondrial-targeted superoxide dismutase mimetic mitoTEMPO, since cytosolic ROS is thought to be largely derived from mitochondria (58). However, neither NAC nor mitoTEMPO improved replication of the double transporter mutant strain under the conditions tested (**S4 Fig**). This observation suggests that as-yet-undefined Mn-dependent processes are required for the cytosolic lifestyle of *Lmo*.

To evaluate the importance of MntH and MntABC over several intracellular replication cycles, we monitored the ability of the transporter single and double mutants to form plaques in L2 fibroblast monolayers. We did not observe any decrease in plaque area for either of the single mutant strains relative to wild type (**Fig 5E**), supporting the notion that either MntH or MntABC is sufficient to facilitate cytosolic replication in this model. While we did observe slight, but statistically significant, increases in plaque area for the single mutant strains in some cases (Δ*mntH* in **Fig 5E** and Δ*mntA* e.v. in **Fig 5F**), these increases were not observed across all experiments (**Fig 5E**-**G**) and thus may not be biologically relevant. Additionally, while others have reported reduced plaque numbers for a Δ*mntA* strain during infection of HT-29 epithelial cells (47), we did not observe substantive differences in plaque number between wild type and the Δ*mntA* strain in our fibroblast model (54 vs. 48 plaques, respectively, across three replicate experiments). Strikingly, however, the double mutant strain was totally unable to form plaques (**Fig 5E**). The plaquing defect of this strain was completely restored upon genetic complementation of either *mntH* or *mntA* (**Fig 5F**-**G**). Cumulatively, these data indicate that the ability to scavenge Mn is essential for the intracellular lifestyle of *Lmo*.

### Mn scavenging is required for virulence of *Lmo*

Finally, we tested the hypothesis that Mn acquisition is required for virulence of *Lmo* using a systemic infection model. C57BL/6 mice were intravenously infected with wild-type, Δ*mntH,* Δ*mntA*, and Δ*mntH* Δ*mntA* strains of *Lmo*. Forty-eight hours post-infection, animals infected with wild-type, Δ*mntH*, or Δ*mntA* strains of *Lmo* lost ∼15% of their body weight (**Fig 6A**), indicating that they are exhibiting signs of listeriosis. Bacterial burdens in the livers and spleens of animals infected with the Δ*mntH* mutant were similar to those infected with wild type, suggesting that MntH is not necessary for systemic listeriosis (**Fig 6B**-**C**). We did observe a modest, trending decrease in bacterial burdens in both the liver and spleen for mice infected with the Δ*mntA* strain, though these values were not statistically significant. Strikingly, the Δ*mntH* Δ*mntA* double mutant was severely attenuated, with a ∼5-log decrease in liver burden and a ∼4-log decrease in spleen burden relative to wild type (**Fig 6B**-**C**). In concordance with this, animals infected with the Δ*mntH* Δ*mntA* strain maintained their body weight over the two days of infection (**Fig 6A**). Altogether, these data are consistent with Mn scavenging via either MntH or MntABC being an essential virulence determinant for *Lmo*.

**Fig 6.**
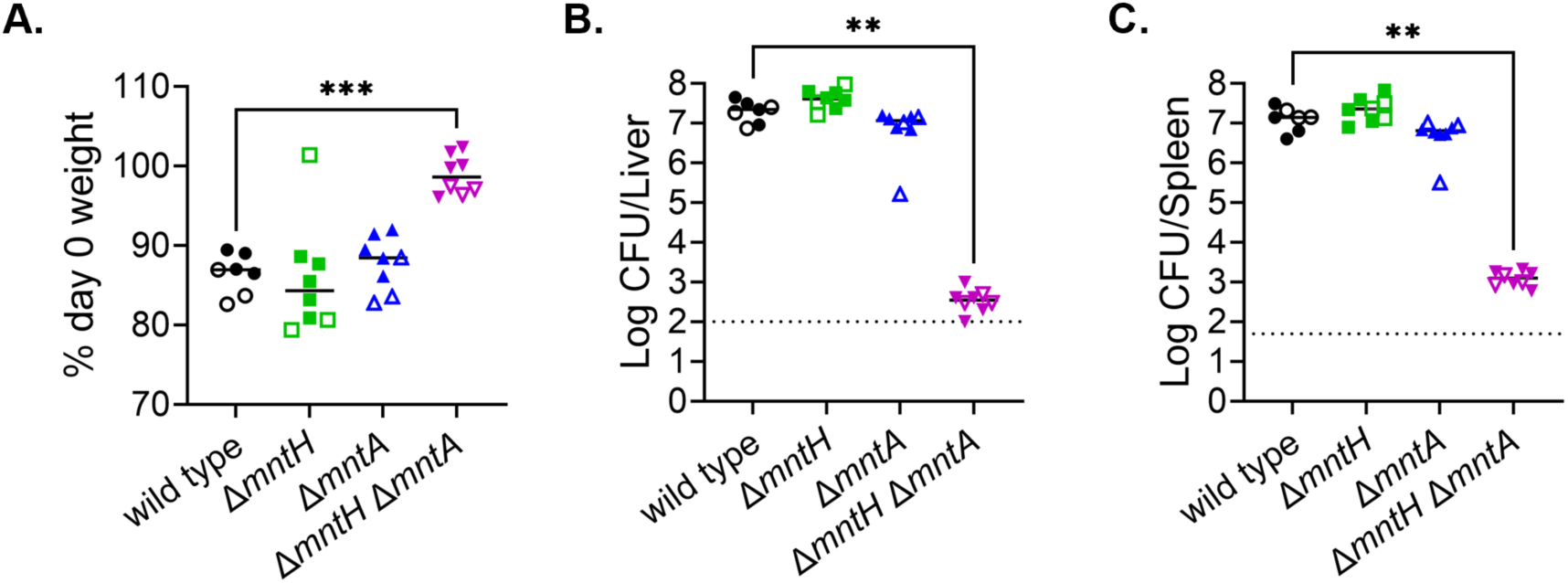
Either MntH or MntABC is required for virulence of *Lmo* in a systemic model of listeriosis. Seven-week-old C57BL/6 mice were infected intravenously with 1 × 10^5^ CFU of the indicated strains of *Lmo*. (A) Endpoint weights of *Lmo*-infected mice are expressed as a percentage of their weight at day 0. Solid bars indicate the median. Statistically significant differences relative to wild type are shown and *** indicates p ≤ 0.001 by one-way ANOVA with Tukey’s multiple comparisons. (B) Livers and (C) spleens were harvested at 48 hr post-infection and bacteria were enumerated by CFU plating. Dotted lines indicate the limit of detection for each organ. Statistically significant differences relative to wild type are shown and ** indicated p ≤ 0.01 by Kruskal-Wallis test with Dunn’s multiple comparisons. Data are pooled from two independent experiments with either male (closed symbols) or female (open symbols) mice. N = 7 for wild type, n = 8 for all other strains.

## Discussion

In this study, we set out to characterize the role of MntH and MntABC homologs in Mn acquisition in *Lmo* and evaluate their contribution to intracellular survival and virulence. Consistent with prior work, we found that neither transporter is essential individually in rich medium under neutral pH conditions (18, 44, 45, 47). However, we found that simultaneous disruption of both loci abrogated growth in a Mn-dependent manner and resulted in reduced intracellular Mn relative to wild type under Mn-limiting conditions. Together these results are consistent with MntH and MntABC functioning as Mn importers in *Lmo*. Further, these data suggest that MntH and MntABC are the only high-affinity Mn transporters in *Lmo* expressed under the conditions tested herein. Then, using the double mutant strain that is unable to scavenge Mn, we were able to demonstrate that high-affinity Mn import is required for adaptation of *Lmo* to its cytosolic niche and for virulence.

While our data suggest that MntH and MntABC are high-affinity Mn transporters and are likely to be the only such transporters in *Lmo*, our observation that the Δ*mntH* Δ*mntA* strain was unable to grow on BHI agar (**Fig 1A**) was still fairly striking given that we did not deplete this rich medium of its normal Mn content and that non-cognate transporters should be sufficient to support growth under Mn-replete conditions. Other groups have measured the Mn concentration in BHI to be in the range of ∼0.3-26 µM (24, 52, 59), though values are expected to be inherently variable between lots and preparations due to the complex nature of BHI medium. Similar observations were made for strains of *Streptococcus mutans* and *E. faecalis* lacking *mntH* and *mntA* homologs in these organisms (24, 52), though double transporter mutants in other species seem to be able to grow in BHI, such as *S. aureus* (20) and *N. gonorrhoeae* (59). These observations likely reflect differences in relative metabolic and nutrient demands of these organisms. Our data also indicate that either MntH or MntABC is able to support growth of *Lmo* in BHI, although the Δ*mntA* mutant (but not Δ*mntH*) exhibited a slight but reproducible defect in BHI liquid culture that could be genetically complemented (**Fig 1B, S1D Fig**) and had lower intracellular Mn under limiting conditions by ICP-MS (**Fig 2B**). Together, this suggests that MntABC may be the primary Mn transporter under these growth conditions. Wu *et al*. reported that disruption of *mntH* alone is sufficient to impair growth in a Mn-dependent manner under acidic conditions (18), consistent with the characterization of MntH homologs as Mn^2+^/H^+^ symporters. Thus, it seems that MntABC and MntH may differentially contribute to Mn acquisition under different environmental conditions. Whether MntABC and MntH have promiscuity for other divalent metals remains unclear. Previous work suggested that MntH may play some role in Zn acquisition (18), and we observed modest growth improvement for strains lacking MntA in BHI supplemented with Fe (**Fig 1C, S1B Fig**). However, only Mn supplementation fully rescues the growth defect of the Δ*mntH* Δ*mntA* strain in BHI (**Fig 1C**) or a *mntH* null strain grown at low pH (18), leading us, and others, to conclude that the primary physiological role of MntH and MntABC is Mn acquisition.

As additional evidence that MntH and MntABC function as Mn transporters in *Lmo*, we demonstrated that they are transcriptionally regulated in response to Mn availability (**Fig 3**). This is consistent with the regulatory scheme defined for homologous transporters in other organisms in which high intracellular Mn levels lead to repression of *mntH* and *mntABC* transcription by the Mn-sensitive transcriptional regulator MntR to prevent toxic accumulation of Mn (20, 49–53). When intracellular Mn levels drop, transporter expression is derepressed to allow for scavenging. Here, we observed 12- and 60-fold induction of *mntH* and *mntA* respectively under Mn-limiting growth conditions (0.1 µM Mn) relative to Mn-replete conditions (50 µM Mn). Derepression of Mn transporter loci in *B. subtilis* and *S. aureus*, which is dependent on MntR, was similarly observed in Mn concentrations below 0.1 µM Mn (20, 49–51). This, and the fact that *Lmo* encodes an *mntR* homolog in its genome (18, 60), suggests that *Lmo* may similarly utilize MntR for Mn-dependent regulation of its Mn importers. Indeed, Wu *et al*., identified potential MntR binding motifs upstream of both *mntH* and the *mntABC* operon (18). Beyond this, the role of MntR in regulating Mn homeostasis has not yet been examined in *Lmo* but will be explored in greater detail in future studies.

We hypothesized that one potential explanation for the high demand of *Lmo* on Mn was the fact that *Lmo* encodes only a single Mn-dependent SOD (54–56) and that Mn also reduces ROS stress by alleviating the reliance on Fe as a cofactor (7, 8). Further, disruption of Mn scavenging in other organisms has been linked to increased sensitivity to oxidative stress (11, 17, 20, 24, 51, 52, 59, 61). In line with this, we found that the *Lmo* Δ*mntH* Δ*mntA* strain is more sensitive to the intracellular superoxide-generating agent PQ *in vitro* and that anaerobic conditions partially restored growth of this strain on BHI agar (**Fig 4**). Together, this suggests that Mn acquisition is important for *Lmo’*s ability to mitigate oxidative stress. However, we did not detect any significant impairment of SOD activity in *Lmo* lysates during Mn-limited growth even during PQ stress (**S3B Fig**), which is somewhat incongruous with the presumed dependence of *Lmo* on Mn for SOD activity and observed PQ sensitivity. While we did observe differences in intracellular Mn concentration in wild-type vs. Δ*mntH* Δ*mntA Lmo* grown under the conditions used for the SOD activity assay, this did not manifest in significantly altered growth kinetics (**Fig 2B, S2C Fig**). Thus, we hypothesize that under our assay conditions the Δ*mntH* Δ*mntA* may not be experiencing a sufficient level of Mn deprivation to see significant differences in SOD activity. Alternatively, it may be that there is a lower limit for SOD activity necessary to sustain growth during Mn limitation that is precluding us from observing differences between strains.

Using an *ex vivo* model of infection, we found that that Δ*mntH* Δ*mntA* strain is impaired in its ability to replicate within macrophages (**Fig 5A**). Given that our *Lmo* strains were grown in Mn-replete medium prior to infection of non-activated macrophages, and that the double mutant was still able to replicate somewhat between 2-5 hr post-infection in BMDMs, which only occurs after *Lmo* escapes the vacuole (62), our data point to a specific defect in adaptation once *Lmo* reaches the cytosol. Supplementation of the cell culture medium with exogenous Mn, which has previously been shown to increase intracellular labile pools of Mn in other mammalian cell lines (63), improved intracellular replication of the Δ*mntH* Δ*mntA* strain (**Fig 5D**). Together, these findings are consistent with *Lmo* being challenged by Mn limitation in the macrophage cytosol. Earlier work found that a Δ*mntA* mutant of *Bacillus anthracis,* the spore-forming causative agent of anthrax that germinates within alveolar macrophages, was delayed in growth within macrophages *ex vivo* in a Mn-dependent manner (26). While vegetative *B. anthracis* do escape the phagolysosome and reach the macrophage cytosol (64), it is not clear whether the delayed growth of the Δ*mntA* strain in this study was due to Mn limitation within the cytosol specifically. However, studies using both *Shigella flexneri,* a cytosolic pathogen that replicates within epithelial cells, and *S.* Typhimurium, which can access and replicate in the cytosol of epithelial cells at low frequency (65), provide evidence that the cytosol of epithelial cells is Mn-limiting for these organisms. High-affinity Mn transporters in both *S. flexneri* and *S.* Typhimurium are specifically upregulated during cytosolic growth (36, 61, 66). Additionally, simultaneous deletion of the MntH and MntABC homologs in *S.* Typhimurium reduced the proportion of bacteria in the cytosol relative to the *Salmonella*-containing vacuole, consistent with Mn limitation restricting growth specifically in the cytosol (36). Loss of these transporters in *S. flexneri*, however, had no impact on plaque formation in an epithelial cell line, but did result in decreased survival in activated macrophages (61). Thus, our findings add to the growing body of evidence that Mn is limiting to pathogens in the cytosol of various cell types and establish that adaptation to Mn limitation is a requirement for survival of *Lmo*, a professional cytosolic pathogen, within its primary replicative niche. Future studies employing the use of Mn transporter transcriptional reporters will help further characterize the extent to which *Lmo* experiences Mn limitation in the cytosol and during infection more broadly.

Despite the increased sensitivity of the Δ*mntH* Δ*mntA* strain to oxidative stress *in vitro*, we found that we were unable to rescue the intracellular growth defect of the double mutant strain in macrophages with either of the ROS-mitigating agents NAC or mitoTEMPO (**S4 Fig**). Thus, we hypothesize that while ROS stress is likely relevant to survival and virulence of *Lmo* in the larger context of systemic infection, it may not be a significant stress experienced by *Lmo* in the cytosol of non-activated macrophages. In line with this, while deletion of *sod* highly attenuates virulence in animal models and was found to increase killing of *Lmo* in IFN-γ-activated macrophages, it has no effect on replication of *Lmo* within non-activated macrophages (67). In the absence of the phagocytic oxidative burst, it appears that superoxide stress is not restrictive to vacuolar escape or cytosolic replication of *Lmo.* Additionally, while *Lmo*’s lone catalase, Kat, is required for normal aerobic growth *in vitro*, it is dispensable for intracellular survival in non-activated BMDMs (68), as are the multiple peroxidase enzymes encoded by *Lmo* (69). Thus, it seems likely that impairment of intracellular replication within BMDMs we observed for the Δ*mntH* Δ*mntA* strain was instead due to dysregulation of another Mn-dependent cellular process that is critical for adaptation to the cytosol beyond ROS stress response. Indeed, multiple pathways with demonstrated roles in cytosolic adaptation of *Lmo*, such as glycolysis (70, 71), menaquinone biosynthesis (72), and maintenance of cell envelope homeostasis (33, 73), are predicted to require Mn-dependent enzymes. These include GpmI, a predicted Mn-dependent phosphoglycerate mutase; MenD, the first dedicated enzyme in menaquinone biosynthesis; PrpC, the cognate phosphatase for the cell envelope stress-sensing kinase, PrkA; and D-alanine-D-alanine ligase (Ddl) (UniProt). Likely, Mn plays multifaceted roles in cytosolic adaptation of *Lmo* that will require further study to uncover.

Ultimately, our *in vivo* infection studies revealed a pronounced attenuation of bacterial burdens in the liver and spleen for mice infected with the Δ*mntH* Δ*mntA* strain (**Fig 6B**-**C**) indicating that Mn scavenging is essential for virulence of *Lmo*. Both *mntH* and *mntABC* were identified in a transcriptomics screen as loci that were upregulated in *Lmo* isolated from the spleen of infected mice (48), consistent with the idea that *Lmo* experiences Mn starvation *in vivo*. Our use of an intravenous model of infection allowed us to focus our attention on the systemic phase of *Lmo* infection; however, this approach inherently bypasses the natural route of *Listeria* infection, *i.e.*, via consumption of contaminated food and transversal of the epithelial barrier. Thus, future studies that address the role of Mn scavenging for fitness in the gastrointestinal tract are warranted, especially given that competition with other bacterial species for Mn has been demonstrated to restrict *Lmo* replication, at least in food cultures (42). Furthermore, Norberg *et al.* recently found that the host divalent metal transporter Nramp2 is expressed in intestinal epithelial cells and contributes to Mn restriction within the *Salmonella*-containing vacuole in this cell type (74), providing precedent for nutritional immunity employed by a non-immune cell. Whether *Lmo* similarly experiences Mn starvation (mediated by Nramp2 or not) during replication in intestinal epithelial cells may be of interest. Use of an oral infection model could also help to explore distinct functions of MntH and MntA that have been proposed by earlier studies, such as the role of MntH in acid tolerance (18), which may be relevant to survival in the gastrointestinal tract, and the potential role of MntA in invasion, which is critical for virulence in this model. Earlier work showed reduced intracellular burdens of a Δ*mntA* strain at early timepoints in *ex vivo* infections of epithelial and hepatocyte cell lines and reduced plaque number on HT-29 cells (44, 47), potentially consistent with an invasion defect in this strain. While we did not observe any major reduction in plaque number or size for the *mntA* deletion strain relative to wild type in our fibroblast infection experiments that might indicate an invasion defect (**Fig 5E**), we did not formally investigate the role of MntA in invasion in this work. Additional *ex vivo* and *in vivo* studies may be warranted to address this more directly, particularly given that genetic complementation of the *mntA* deletion allele was not demonstrated in earlier work in which invasion defects were reported (44, 47).

Taken together, this work provides evidence that MntH and MntABC are the sole high-affinity Mn transporters in *Lmo* and are necessary for adaptation of this pathogen to its cytosolic replicative niche and ultimately virulence in an animal model. This suggests that *Lmo* experiences Mn limitation within the cytosol, but whether specific nutritional immunity strategies exist that contribute to Mn limitation within this niche during infection remains unknown. Future exploration of changes in both Mn mobilization and expression or localization of host Mn transporters in response to *Lmo* infection will help address this possibility.

## Material and methods

### Ethics statement

All experiments involving animals were approved by the Institutional Animal Care and Use Committee of the Medical College of Wisconsin.

### Bacterial strains, cloning, and growth conditions

*Lmo* was routinely grown at 37°C in complete Listeria Synthetic Medium (LSM, (43)) containing 50 µM MnSO_4_ (Sigma-Aldrich) or on brain heart infusion (BHI; VWR) + 1.5% agar plates (supplemented with 50 µM MnSO_4_ for the Δ*mntH* Δ*mntA* strain). Strains were frozen down in BHI + 40% glycerol. *Escherichia coli* strains were routinely grown at 37°C in LB or on LB + 1.5% agar plates and frozen in LB + 40% glycerol. Antibiotics were used at the following concentrations: 100 µg/mL ampicillin, 10 µg/mL chloramphenicol, 30 µg/mL kanamycin, 200 µg/mL streptomycin, 20 µg/mL nalidixic acid, 2 µg/mL erythromycin and 50 µg/mL gentamicin. Primers (**Table 1**), plasmids (**Table 2**) and strains (**Table 3**) used in this study are listed below. *E. coli* XL1-Blue was used as a routine cloning strain. Deletions were generated by allelic exchange using *p-*chloro-phenylalanine counterselection as described previously (75), leaving the first and last 4-8 codons in frame. The pLIM1-Δ*mntH* and pLIM1-Δ*mntA* plasmids were generated by Gibson Assembly using pLIM1 (from Arne Rietsch, Case Western Reserve) linearized with BamHI and two ∼1 kb PCR-generated insert fragments amplified from *Lmo* 10403S genomic DNA each. CAM90 and CAM91 were used to amplify the region upstream of *mntH* while CAM92 and CAM93 were used to amplify the *mntH* downstream fragment. CAM123 and CAM124 were used to amplify the *mntA* upstream fragment while CAM125 and CAM99 were used to generate the *mntA* downstream fragment. The pIMK2-*mntA* plasmid was generated by Gibson assembly using pIMK2 linearized with BamHI and SalI and a PCR fragment of the *mntA* gene amplified from 10403S gDNA using primers CAM157 and CAM158. pPL2e-P*_mntH_*-*mntH* was also generated by Gibson assembly using pPL2e linearized with SmaI and EagI and a PCR fragment amplified with CAM166 and CAM168 which included the *mntH* gene as well as the 112 bp immediately preceding the gene (predicted to include the native promoter based on BPROM analysis) and 86 bp downstream of the gene. All plasmid inserts were verified by Sanger sequencing. Plasmids were shuttled into *Lmo* via conjugation using *E. coli* strains SM10 (for pLIM1 and pPL2e) or S17 (for pIMK2).

**Table 1.**
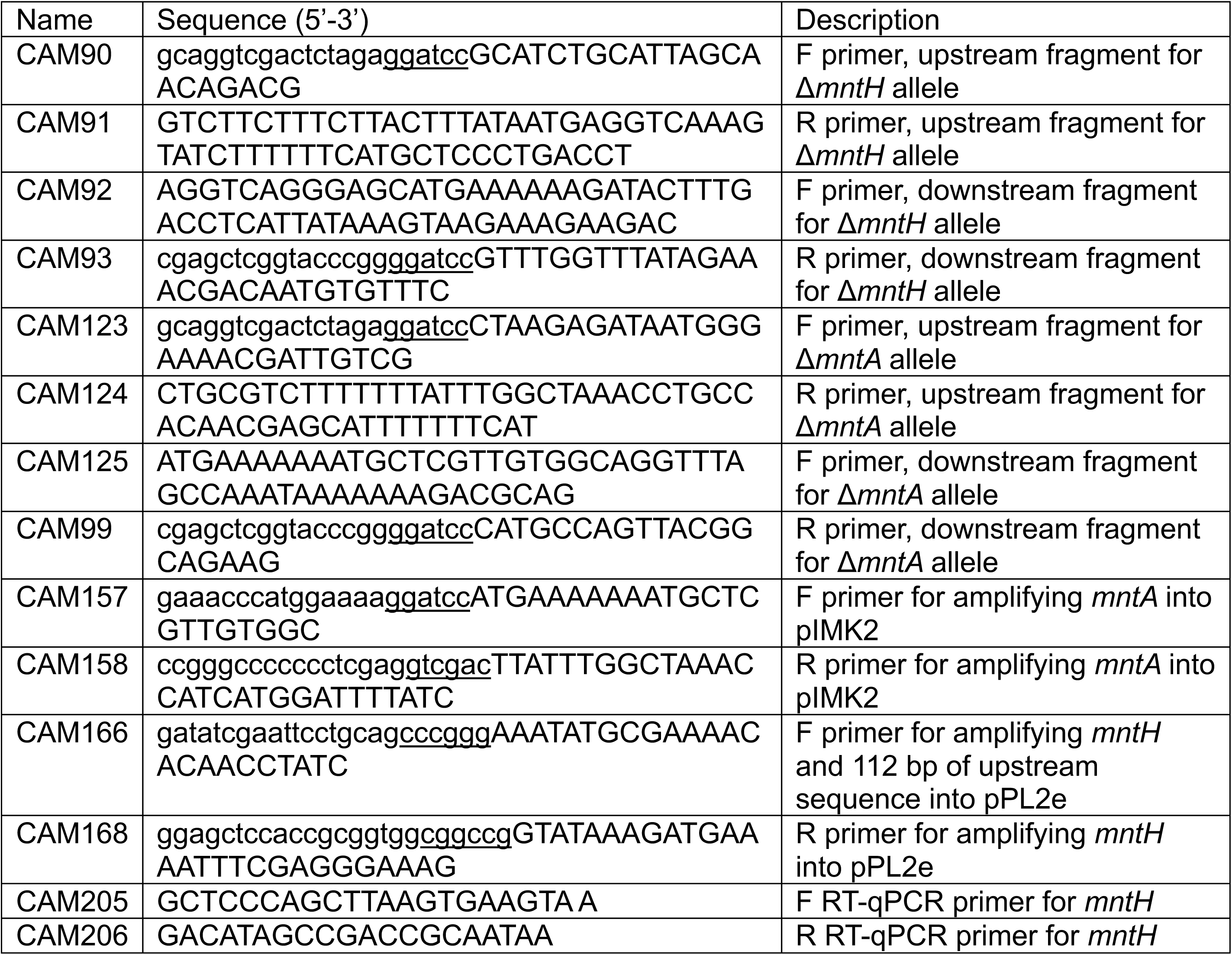
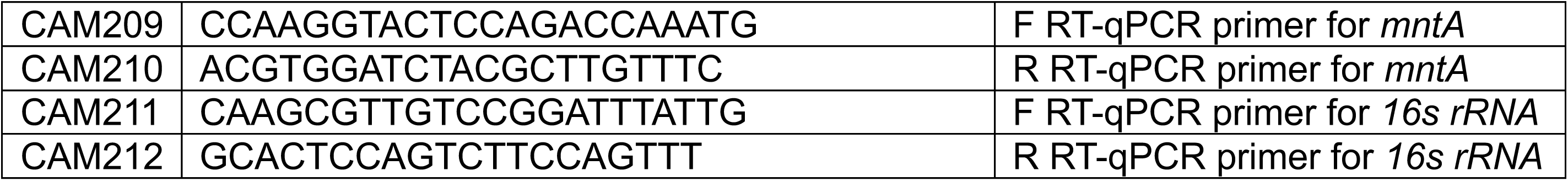
Primers used in this study.

**Table 2.**
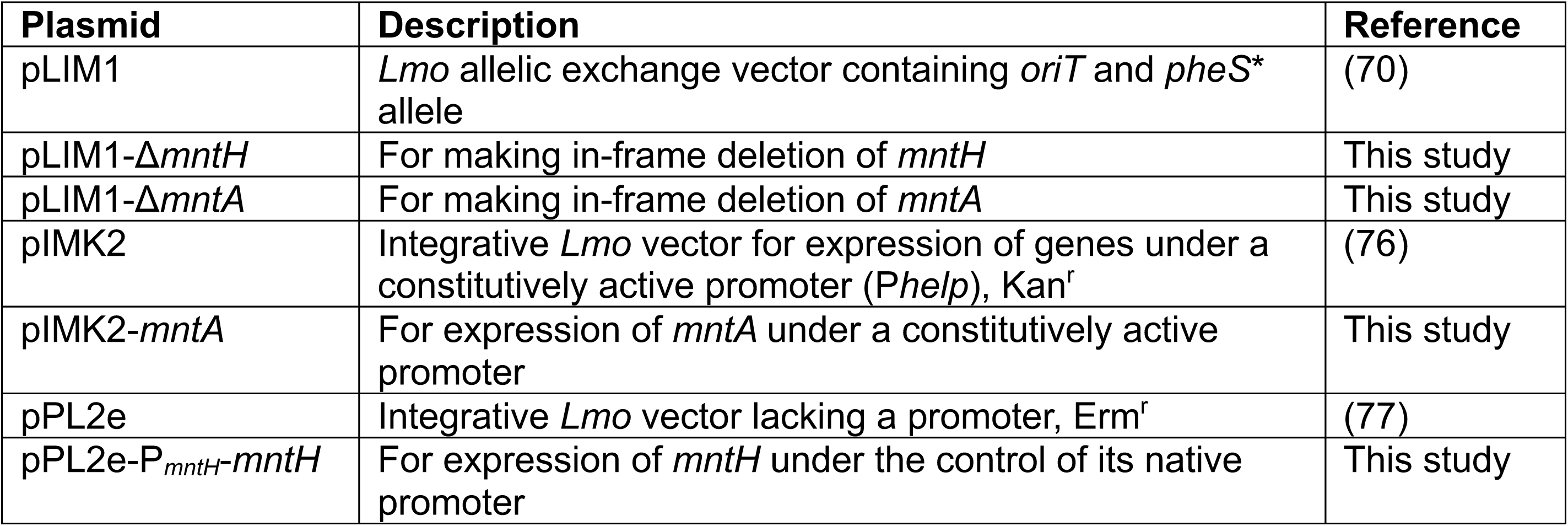
Plasmids used in this study.

**Table 3.** Strains used in this study.

| *Lmo* strains | Strain number | Genotype/Description | Reference |
| --- | --- | --- | --- |
| wild type | JLK1 | *Lmo* strain 10403S | (78) |
| $\Delta mntH$ | JLK1622 | In-frame deletion of *mntH* via pLIM1-$\Delta mntH$ | This study |
| $\Delta mntA$ | JLK1860 | In-frame deletion of *mntA* via pLIM1-$\Delta mntA$ | This study |
| $\Delta mntH \Delta mntA$ | JLK1859 | In-frame deletion of *mntA* via pLIM1-$\Delta mntA$ in the $\Delta mntH$ background | This study |
| wild type pIMK2 | CM373 | wild type integrated with pIMK2 | This study |
| $\Delta mntH$ pIMK2 | JLK2057 | $\Delta mntH$ integrated with pIMK2 | This study |
| $\Delta mntH \Delta mntA$ pIMK2 | JLK2060 | $\Delta mntH \Delta mntA$ integrated with pIMK2 | This study |
| $\Delta mntH \Delta mntA$ pIMK2-*mntA* | JLK2061 | $\Delta mntH \Delta mntA$ integrated with pIMK2-*mntA* | This study |
| wild type pPL2e | JLK2062 | wild type integrated with pPL2e | This study |
| $\Delta mntA$ pPL2e | JLK2064 | $\Delta mntA$ integrated with pPL2e | This study |
| $\Delta mntH \Delta mntA$ pPL2e | JLK2065 | $\Delta mntH \Delta mntA$ integrated with pPL2e | This study |
| $\Delta mntH \Delta mntA$ pPL2e-P<sub>*mntH*</sub>-*mntH* | JLK2066 | $\Delta mntH \Delta mntA$ integrated with pPL2e-P<sub>*mntH*</sub>-*mntH* | This study |

### Growth assays

*Lmo* strains were grown overnight (18-20 hr) in 2 mL of LSM containing 50 µM MnSO_4_ at 37°C with shaking. Overnight cultures were then normalized to OD_600_ = 1.0 and washed once with PBS. For spot plate assays, the washed cells were resuspended in BHI then serially diluted. 4 µL of each dilution was spotted on either BHI agar or BHI agar with 50 µM supplemental MnSO_4_ then plates were incubated at 37°C for ∼24 hr. For anaerobic growth, plates were placed in a BD GasPak EZ Anaerobe Pouch System with an indicator strip. Plate images were captured using an Azure 300 Imaging System (Azure Biosystems, Inc., Dublin, CA). For BHI growth curves, washed cells were resuspended in BHI then diluted 1:50 into 100 µL of BHI in flat-bottom 96-well plates. When indicated, MnSO_4_, FeCl_2_, and ZnSO_4_ were added to the medium at the concentrations denoted in the figure legends. OD_600_ was measured every 15 min during growth at 37°C on either a BioTek Synergy H1 or BioTek Epoch 2 microplate reader (Agilent BioTech, Santa Clara, CA). Plates were subject to continuous double orbital shaking between measurements. For LSM growth curves, washed cells were resuspended in LSM lacking MnSO_4_ then diluted 1:50 into 100 µL LSM + 0.1% Tween80 in round-bottom 96-well microplates (Thermo Scientific). OD_600_ measurements were made as described above for BHI growth curves. When indicated, paraquat (PQ; Sigma-Aldrich) was added to the medium at 1 mM.

### *Lmo* harvest for ICP analysis

To measure intracellular metal content, *Lmo* strains were grown overnight (18-20 hr) in 10 mL of LSM containing 50 µM MnSO_4_ at 37°C with shaking in 50 mL conical tubes. Overnight cultures were normalized to OD_600_ = 1.0, washed twice with PBS and resuspended in LSM lacking MnSO_4_. In duplicate, cells were then diluted 1:6 in 15 mL final volume of LSM containing either 50 or 0.1 µM MnSO_4_ in metal-free 50 mL conical tubes (VWR) then incubated at 37°C with shaking. Cultures were harvested at an OD_600_ = 0.70-0.80, and duplicate cultures were combined and subjected to centrifugation to form cell pellets. Pellets were washed twice with 0.1 M EDTA followed by two washes with ultrapure H_2_O. Cells were then plated for CFU enumeration on BHI agar (with 50 µM MnSO_4_ added for the Δ*mntH* Δ*mntA* strain) and the remaining pellet was dried on a heat block at 94°C overnight. Replicates were collected across three separate experiments.

### Reagents, standard, and sample preparation for ICP analysis

All standards, blanks, and samples were prepared using sub-boiling point distilled (Savillex DST-1000, Eden Prairie, MN, USA) trace metal grade nitric acid (70%, Fisher chemical, Cat# A509P212), ultrapure water (18.2 MΩ·cm @ 25°C), and metal free polypropylene conical tubes (15 and 50 mL, Labcon, Petaluma, CA, USA) and measured gravimetrically. Multi-element custom standard containing 1000 µg/mL of As, B, Ca, Cd, Co, Cr, Cu, Fe, K, Mg, Mn, Mo, Na, Ni, Pb, S, Se, V, Zn and a single element 1000 µg/mL P standard (Inorganic Ventures; Christiansburg, VA) were diluted to 100, 50, 25, 12.5 6.25, 3.125, and 0 ng/g in 3% (v/v) HNO_3_ in ultrapure deionized. Samples preparation was carried out by adding 400 µL of sample followed by addition of 300 µL of 70% HNO_3_ and incubated at 65°C for 2 hours followed by addition of 9.3 mL of de-ionized H_2_O for a resulting solution of 3% (v/v) HNO_3_.

### ICP-QQQ-MS Analysis

ICP-MS samples, standards, and blanks were analyzed using an Agilent 8900 Triple Quadrupole ICP-MS (Agilent, Santa Clara, CA, USA) equipped with the Agilent SPS 4 Autosampler, integrated sample introduction system (ISiS), x-lens, and Micromist nebulizer. Daily tuning of the instrument was accomplished using the manufacturer supplied tuning solution containing Li, Co, Y, Ce, and Tl. Global tune optimization was based on optimizing intensities for ^7^Li, ^89^Y, and ^205^Tl while minimizing oxides (^140^Ce^16^O/^140^Ce < 1.5%) and doubly charged species (^140^Ce++/^140^Ce+ < 2%). Following global instrument tuning, gas mode tuning in He KED was accomplished using the same manufacturer supplied tuning solution. In KED mode (using 100% UHP He, Airgas), intensities for ^59^Co, ^89^Y, and ^205^Tl were maximized while minimizing oxides (^140^Ce^16^O/^140^Ce < 0.5%) and doubly charged species (^140^Ce++/^140^Ce+ < 1.0%) with short term RSDs < 3.5%. Internal standardization was accomplished inline using the ISIS valve and a 200 ng/g of mixed single element standards multi-element internal standard mix containing Bi, In, ^6^Li, Sc, Tb, Y (IV-ICPMS-71D, Inorganic Ventures, Christiansburg, VA, USA) in 3% (v/v) trace nitric acid in ultrapure water. Acquisition parameters were as follows: gas mode = He KED, Acquisition mode = Spectrum, Replicates = 3, Sweeps/replicate = 30, Stabilization time = 10 s, Scan type = Single quad, RF power = 1550 W, Sampling Depth = 8 mm, Nebulizer gas = 1.09 L/min, Plasma gas = 15 L/min, Auxiliary gas = 0.9 L/min, He flow rate = 5 mL/min, OctP Bias = −18.0 V, Axial acceleration = 1 V, Energy discrimination = 3.0 V. Isotopes selected for analysis were: ^55^Mn, ^56^Fe, and ^66^Zn with ^45^Sc, ^89^Y, and ^115^In selected as internal standards. Standards were fit using linear regression with no weighting with an R^2^ > 0.9999 and %RSE < 5%. Sample sequence was run with a continuing calibration blank (CCB) every 3 samples and continuing calibration verification (CCV) every 30 samples.

### RT-qPCR

For *mntH* and *mntA* transcript analysis, cultures were grown largely as described above for ICP-MS sample preparation but were washed only once in PBS and were diluted 1:6 in LSM containing either 50, 1, or 0.1 µM MnSO_4_. Once at an OD_600_ = 0.60-0.70 cells were harvested with equal volume of an ice-cold 50% ethanol/50% acetone solution, washed with cold water and frozen at −80°C. Thawed pellets were suspended in 10 mM Tris, 1 mM EDTA, 15 mg/mL lysozyme and 250 U/mL mutanolysin (pH = 8.0), then incubated for 20 min at 37°C. RLT buffer (Qiagen, RNeasy Kit) with DTT was added to the cells, which were then vortexed on maximum speed for 30 s in the presence of 0.1 mm zirconia beads. Cell debris was removed by centrifugation, and RNA was purified using the RNeasy Kit (Qiagen, Germantown, MD) per the manufacturer’s instructions. dsDNA was removed and cDNA was synthesized using the Maxima First Strand cDNA Synthesis Kit with dsDNAse (ThermoFisher Scientific, Waltham, MA). qPCR was performed using Power SYBR Green PCR Master Mix (Applied Biosystems^TM^, Waltham, MA) and cycling conditions recommended by the manufacturer on a Bio-Rad CFX Connect Plate reader (Bio-Rad, Hercules, CA). Primer efficiencies were determined using 5-point standard curves of pooled cDNA. Fold-changes of *mntA* and *mntH* transcript were determined by the Pfaffl Method, with *16s rRNA* as the reference gene. Replicate cultures were harvested across three separate experiments and qPCR reactions were set up in triplicate for each sample.

### Intracellular growth assays

Bone marrow-derived macrophages (BMDMs) were isolated as previously described from C57BL/6 mice (79) and were seeded at 4 × 10^6^ cells/60 mm dish with coverslips and allowed to adhere overnight (16-18 hr). *Lmo* strains were grown overnight (18-20 hr) in 3 mL LSM containing 50 µM MnSO_4_ at 30°C on a slant, subjected to centrifugation to form cell pellets, resuspended in PBS at 1 × 10^8^ CFU/mL then used to infect BMDMs at MOI = 0.25. Thirty minutes post-infection, the medium was aspirated and the cells were washed twice with pre-warmed PBS, then fresh medium containing gentamicin was added. At the indicated timepoints, coverslips were removed in triplicate for each infection condition and CFU per coverslip was determined largely as previously described (80). CFU enumeration was done using standard BHI agar plates incubated at 37°C for all strains except Δ*mntH* Δ*mntA*, Δ*mntH* Δ*mntA* pIMK2, Δ*mntH* Δ*mntA* pIMK2-*mntA* and Δ*mntH* Δ*mntA* pPL2e strains which were enumerated on BHI agar containing 50 µM supplemental MnSO_4._ For Mn supplementation experiments, 50 µM MnSO_4_ was added to the cell culture medium at the time of BMDM seeding and in subsequent medium changes at the time of infection and gentamicin treatment. All infections were performed in triplicate and average CFU/coverslip at each timepoint across triplicate experiments are reported.

### Plaquing assays

Plaquing assays were performed using the L2 murine fibroblast cell line (81) largely as previously described (82). L2 cells were seeded at 1.2 × 10^6^ cells/well in a 6-well dish 16-18 hr prior to infection. *Lmo* strains were grown and inocula were prepared as described for intracellular replication assays above prior to infection of L2 monolayers at an MOI = 0.5. One hour post-infection, the cells were washed thrice with pre-warmed PBS, then fresh medium containing 0.7% agarose and gentamicin were added to each well. At ∼48 hr post-infection, additional medium containing agarose and gentamicin was added. At ∼72 hr post-infection, agarose plugs were removed, then cells were stained with 0.3% crystal violet for 10 min and subsequently washed twice with water and allowed to dry. Plates were imaged on an Epson scanner, and plaques were quantified and measured using ImageJ software (version 1.53). Individual plaque sizes were normalized to the average value for wild type across experiments. Data reflect the average of all plaques pooled from three replicate experiments.

### Animal infections

Mouse infections were performed intravenously essentially as previously described (83). *Lmo* strains were grown overnight in 3 mL BHI at 30°C on a slant. 50 µM MnSO_4_ was added to the medium for the Δ*mntH* Δ*mntA* strain only. Overnight cultures were normalized to OD_600_ = 1.0, pelleted, then resuspended in BHI without supplemental Mn before being diluted 1:6 in LSM containing 50 µM MnSO_4_ in 125 mL flasks and incubated at 37°C with shaking for 2 hr (OD_600_ = 0.40-0.60). Cells were then pelleted, resuspended in PBS and adjusted to 5 × 10^5^ CFU/mL. Seven-week-old C57BL/6 mice (Charles River Laboratories) were infected with 1 × 10^5^ CFU of *Lmo* in 200 µL of PBS via tail vein injection. Animal weights were recorded at 0, 24 and 48 hr post-infection. At 48 hr post-infection, animals were euthanized by CO_2_ asphyxiation according to AVMA standards. Spleens and livers were harvested, homogenized in PBS + 0.1% NP-40 and plated for CFU on BHI agar plates containing streptomycin and, in the case of the Δ*mntH* Δ*mntA* strain, 50 µM MnSO_4._ Infections were performed in duplicate, one experiment with all female mice and a second experiment with all male mice. Data was pooled from both experiments, and sex is indicated in the figure with closed (male) and open (female) symbols.

### Statistics

Statistical analysis was performed using GraphPad Prism (version 11.1.0). The particular test used for each figure is indicated in the figure legend.

## Acknowledgments

Elemental analysis was performed at the Michigan State University Quantitative Bio Element Analysis and Mapping (QBEAM) Center.

