## Supplementary figures and images for "Manganese scavenging is required for virulence of the cytosolic pathogen *Listeria monocytogenes*"

### S1 Fig

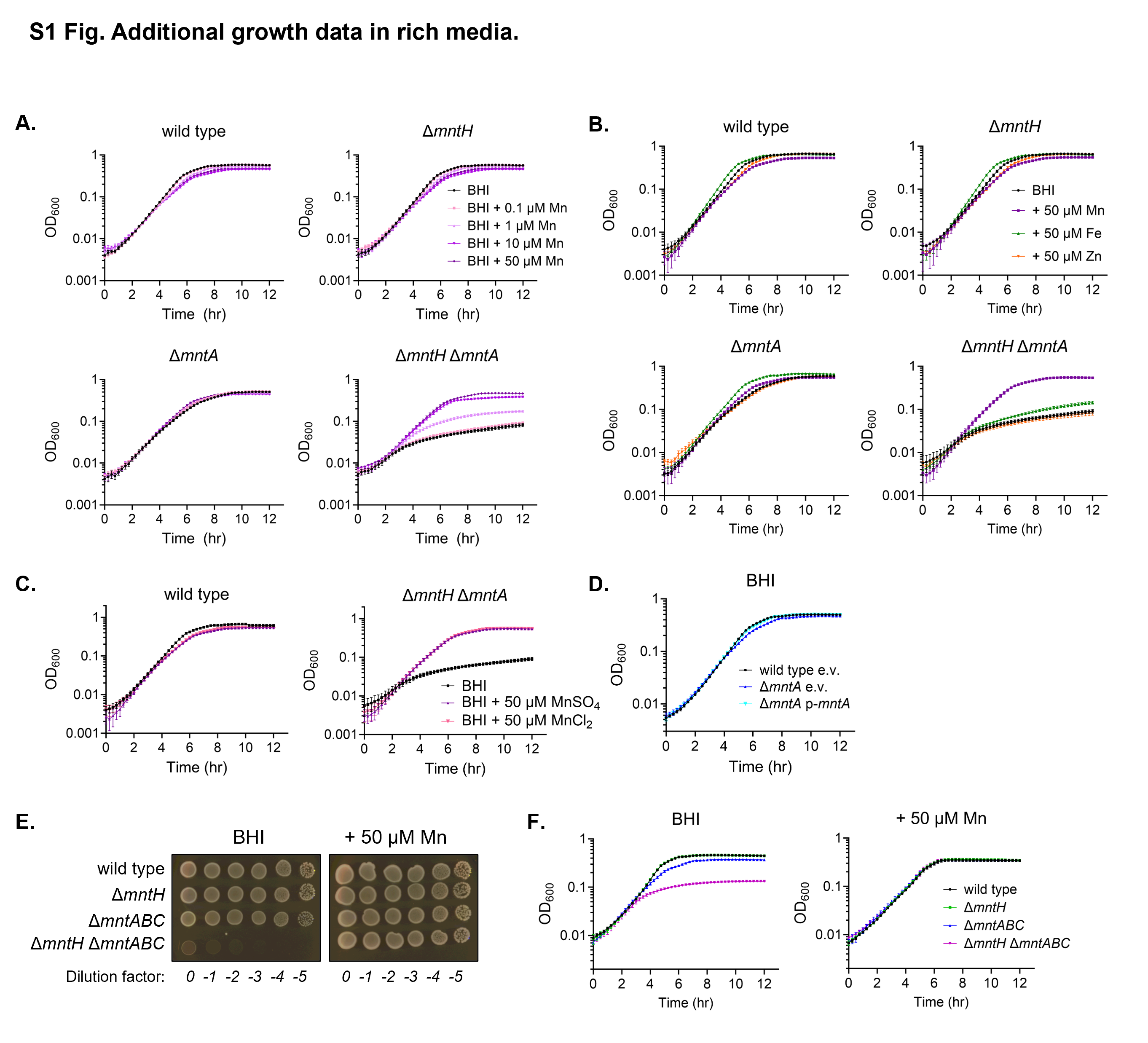

### S2 Fig

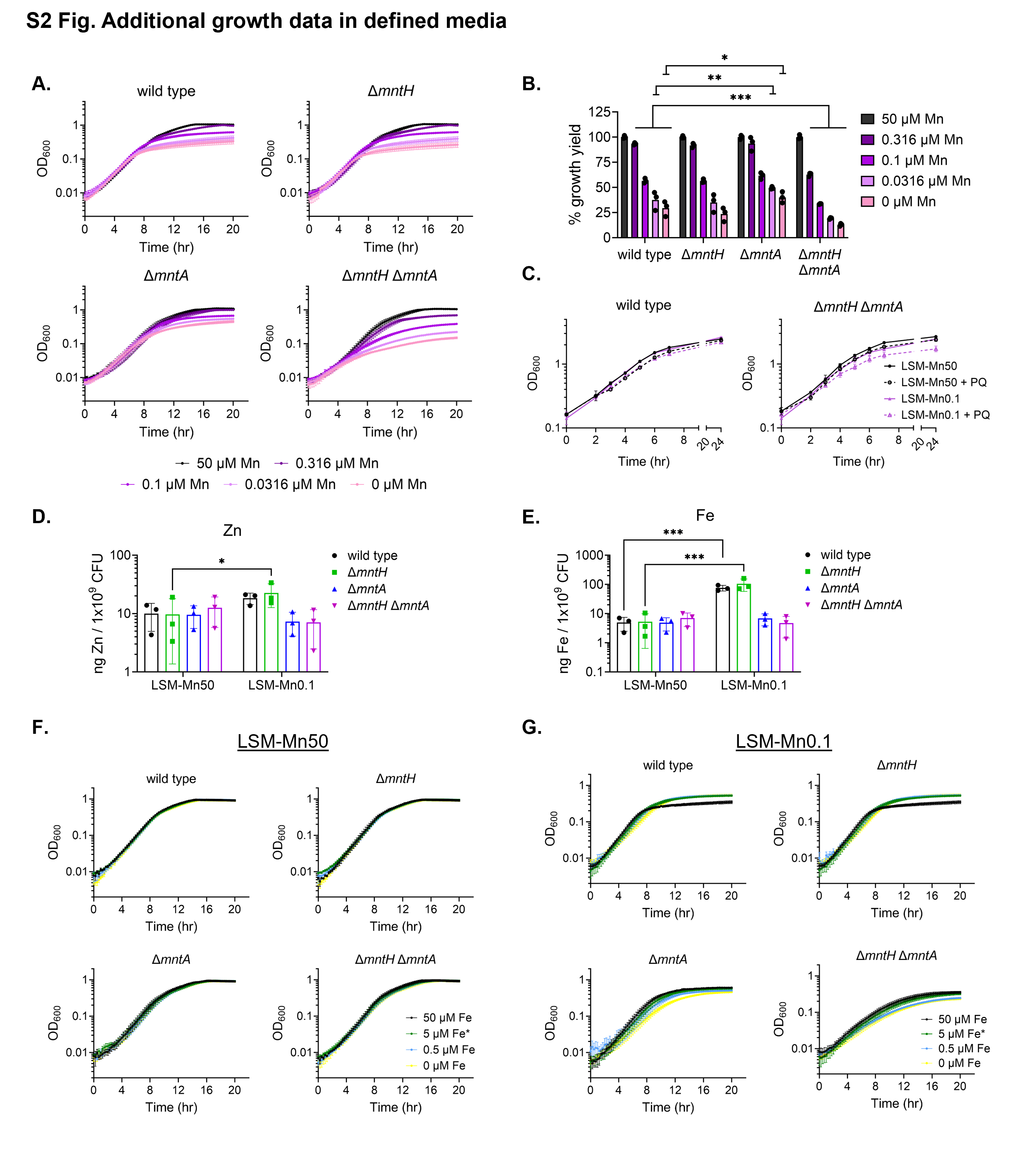

### S3 Fig

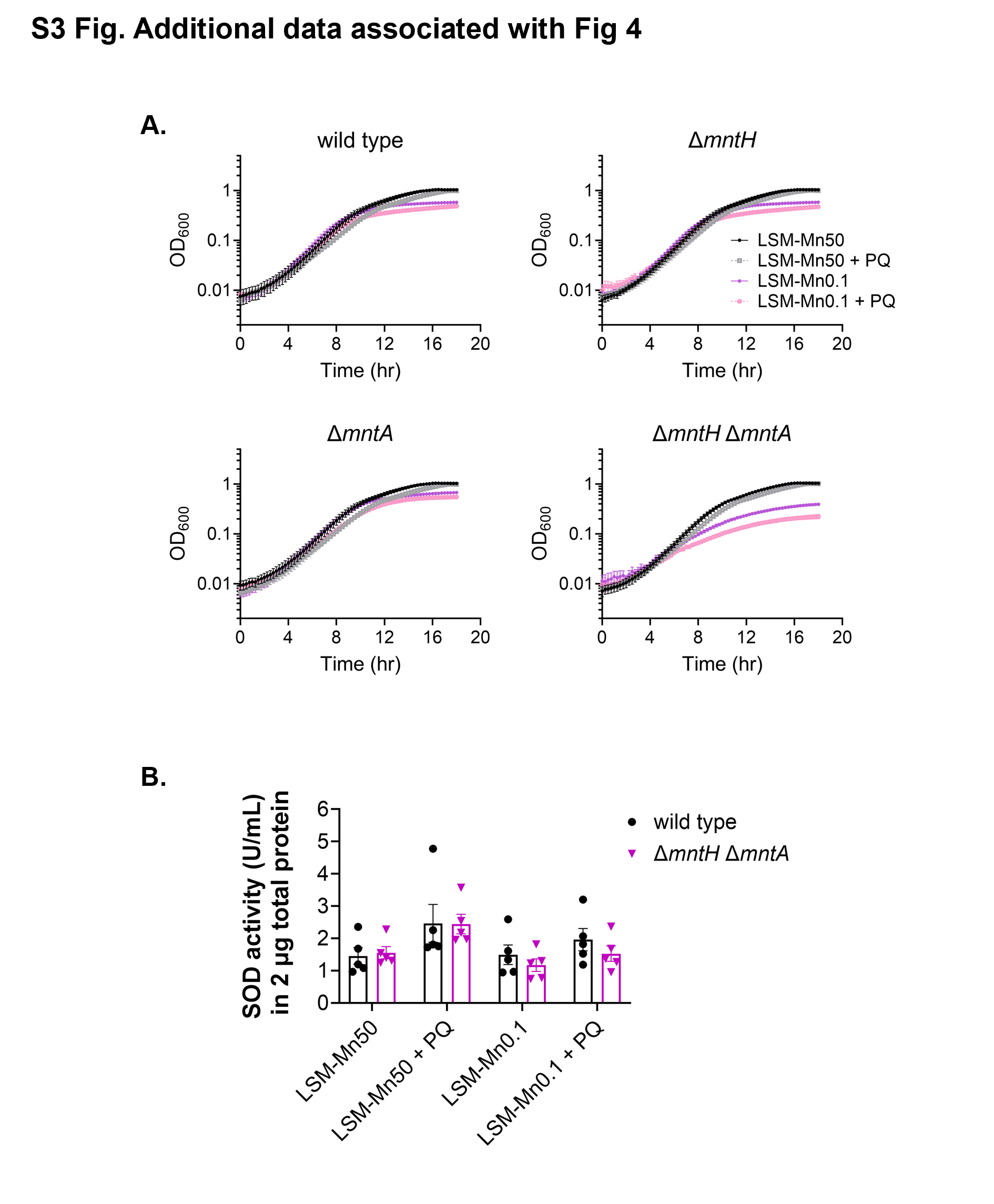

### S4 Fig

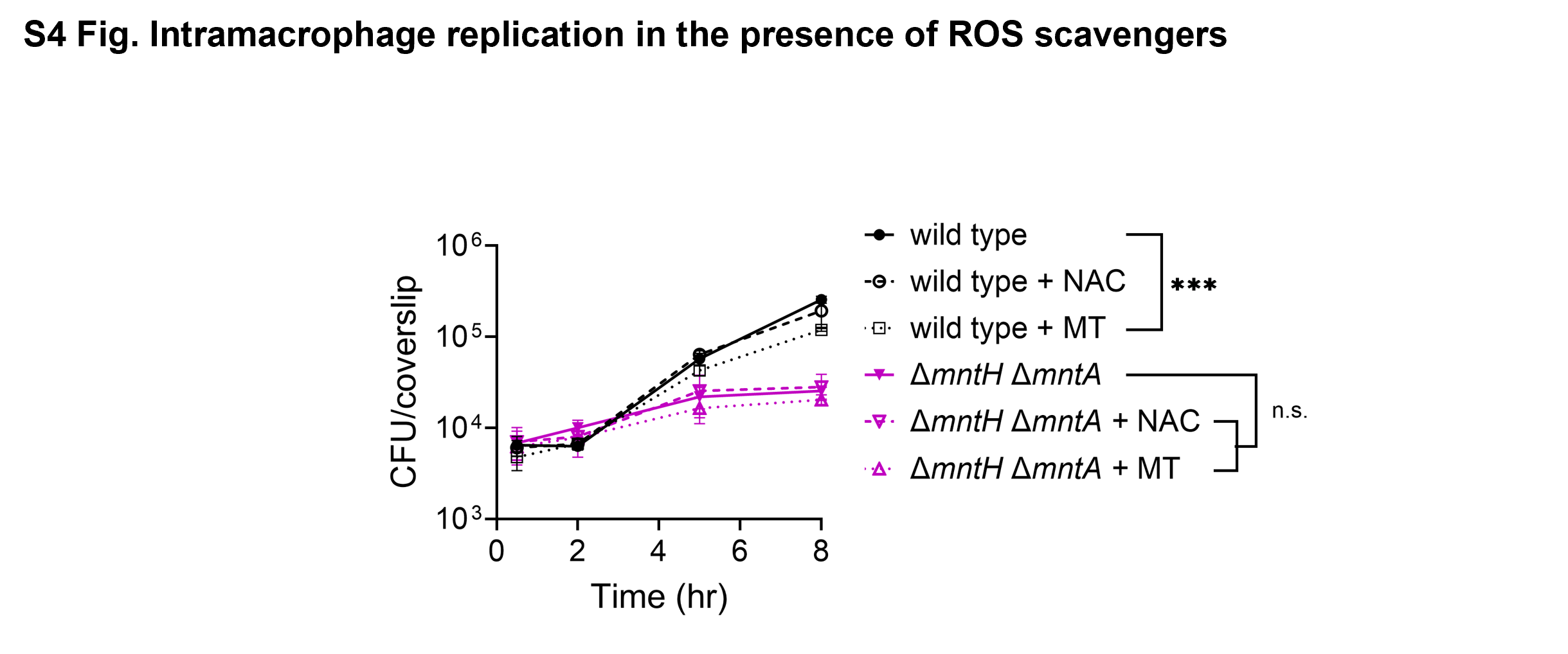
