## Supplementary material for "Manganese scavenging is required for virulence of the cytosolic pathogen *Listeria monocytogenes*": S5 Appendix

**S5 Appendix. Supplemental materials and methods**

**Bacterial strains**

Additional primers (S1 Table), plasmids (S2 Table) and strains (S3 Table) used in supplemental figures in this study are listed below. In-frame deletion of the *mntABC* operon was achieved using pLIM1 as described in the main text. CAM96 and CAM97 were used to amplify a ~1 kb fragment upstream of the operon, while CAM98 and CAM99 were used to amplify a ~1 kb fragment downstream.

**S1 Table. Additional primers used in this study.**

| Name | Sequence (5’-3’) | Description |
| --- | --- | --- |
| CAM96 | gcaggtcgactctagaggatccGGTCCAAACGGCAACAACTTTATC | F primer, upstream fragment for Δ*mntABC* allele |
| CAM97 | CTGCGTCTTTTTTTATTTGGCTAAACCAATTTCCATCTAAACCACCCTTTTCC | R primer, upstream fragment for Δ*mntABC* allele |
| CAM98 | GGAAAAGGGTGGTTTAGATGGAAATTGGTTTAGCCAAATAAAAAAAGACGCAG | F primer, downstream fragment for Δ*mntABC* allele |
| CAM99 | cgagctcggtacccggggatccCATGCCAGTTACGGCAGAAG | R primer, downstream fragment for Δ*mntABC* allele |

**S2 Table. Additional plasmids used in this study**

| **Plasmid** | **Description** | **Reference** |
| --- | --- | --- |
| pLIM1-Δ*mntABC* | For making in-frame deletion of the *mntABC* operon | This study |

**S3 Table. Additional strains used in this study**

| ***L. monocytogenes* strains** | **Strain number** | **Genotype/Description** | **Reference** |
| --- | --- | --- | --- |
| Δ*mntABC* | JLK1624 | In-frame deletion of the *mntABC* operon via pLIM1-Δ*mntABC* | This study |
| Δ*mntH* Δ*mntABC* | JLK1668 | In-frame deletion of the *mntABC* operon via pLIM1-Δ*mntABC* in the Δ*mntH* background | This study |
| Δ*mntA* pIMK2 | JLK2058 | Δ*mntA* integrated with pIMK2 | This study |
| Δ*mntA* pIMK2-*mntA* | JLK2059 | Δ*mntA* integrated with pIMK2-*mntA* | This study |

**SOD activity assay**

*Lmo* strains were grown as described for ICP-MS sample collection, but in the presence or absence of 1 mM paraquat (PQ). Once at an OD_600_ = 0.60-0.80, cells were subjected to centrifugation to form pellets, washed twice with cold PBS, and frozen at -80°C. Thawed pellets were suspended in 50 mM Tris, 150 mM NaCl, 0.1 mM EDTA, 0.1% SDS (pH = 7.4) containing protease-inhibitor cocktail (cOmplete^TM^ Mini; Roche, Basel, Switzerland) and bead beat by vortexing with 0.1 mm zirconia beads for 2 min at maximum speed. Cell debris was removed by centrifugation at 14,000 x g for 10 min, and lysates were normalized by total protein concentration as determined by BCA assay (Pierce^TM^, ThermoScientific^TM^, Waltham, MA). SOD activity in 2 µg of total protein for each sample was determined using the Superoxide Dismutase Activity Assay Kit from Sigma-Aldrich per the manufacturer’s instructions. Absorbance at 450 nm was read on a TECAN Spark microplate reader (Tecan, Männedorf, Switzerland). Samples were collected over 5 replicate experiments and were measured in technical triplicate.

**Intracellular replication assays**

Assays were performed as described in the main text. For supplementation with N-acetylcysteine (NAC; ThermoScientific Chemicals) and mitoTEMPO (MT; Sigma-Aldrich) the compounds were added to the cell culture medium at the time of infection and replaced within the fresh medium during gentamicin treatment at 30 min post-infection. NAC was resuspended in sterile water then neutralized to a pH of 7.4 prior to being added to the cell culture medium at a final concentration of 5 mM. mitoTEMPO was resuspended in DMSO and added at a final concentration of 100 µM.
